# A Logic Approach to Modeling Nomenclatural Change

**DOI:** 10.1101/058834

**Authors:** Nico M. Franz, Chao Zhang, Joohyung Lee

## Abstract

We utilize an Answer Set Programming (ASP) approach to show that the principles of nomenclature are tractable in computational logic. To this end we design a hypothetical, 20 nomenclatural taxon use case with starting conditions that embody several overarching principles of the International Code of Zoological Nomenclature; including Binomial Nomenclature, Priority, Coordination, Homonymy, Typification, and the structural requirement of Gender Agreement. The use case ending conditions are triggered by the reinterpretation of the diagnostic features of one of 12 type specimens anchoring the corresponding species-level names. Permutations of this child-to-parent reassignment action lead to 36 alternative scenarios, where each scenario requires 1-14 logically contingent nomenclatural emendations. We show that an ASP transition system approach can correctly infer the Code-mandated changes for each scenario, and visually output the ending conditions. The results provide a foundation for further developing logic-based nomenclatural change optimization and compliance verification services, which could be applied in globally coordinated nomenclatural registries. More generally, logic explorations of nomenclatural and taxonomic change scenarios provide a novel means of assessing design biases inherent in the principles of nomenclature, and thus may inform the design of future, big data-compatible identifier systems for systematic products that recognize and mitigate these constraints.

---

The origins of modern biological nomenclature trace back to the age of Renaissance (Minelli 2003), culminating in the foundational works of Linnaeus (1753, 1758). The Linnaean rules were subsequently embedded, expanded, and refined in the *Codes* (e.g., ICZN 1999; McNeill et al.2012),thus having remained in use for more than 250 years (Schuh 2003; Dayrat 2010). In spite of such remarkable persistence, the merits of Linnaean nomenclature are continuously under scrutiny (e.g., Bryant and Cantino 2002; Ereshefsky 2007; Dayrat 2010; Dubois 2011; Vences et al. 2013; Remsen 2016). Indeed, the demands on any system for identifying the diversity of life are not trivial, and may include: precise and reliable connections to physical vouchers; congruence with central tenets in evolutionary and systematic theory; responsiveness to changing taxonomic and phylogenetic perspectives; proper attribution of authorship - original or revisionary; stability in usage across spatial and temporal dimensions; transparency in application; support for community-sanctioned rules and updates that accommodate new naming challenges; and facilitation of everyday use among human speakers and general alignment with evolutionarily constrained human cognitive strengths (Atran 1998; Sterner and Franz 2016). No system of nomenclature can fulfill all of these and other potentially conflicting criteria to the highest degree.

Here we introduce a novel element into the longstanding discourse about the Linnaean nomenclature, i.e., its relationship to computational logic. Methods of knowledge representation and reasoning (Brachman and Levesque 2004) are on the rise in biomedical and evolutionary disciplines (Smith et al. 2007; Franz and Thau 2010; Panahiazar et al. 2013; Deans et al. 2015; Thessen et al. 2015). But this dynamic has not yet permeated the realm of nomenclature (though see Sereno 2005; Tuominen et al. 2011; Chawuthai et al. 2013; Dmitriev and Yoder 2016).

We can conceive of at least four reasons why a better integration of nomenclatural practices with logic representation and reasoning is desirable. (1) The language controlled by the rules of nomenclature shapes the ways in which humans communicate much of our collective knowledge about life (Patterson et al. 2010). Hence an exploration of the extent to which these rules are amenable to logic may be of interest in its own right. (2) In light of the current trend towards ontology-driven representation of biodiversity data, it appears relevant to assess whether nomenclatural entities and relationships can, and should, be integrated with these efforts (e.g., Mungall et al. 2010; Midford et al. 2013, Walls et al. 2014). (3) Taxonomic communities such as zoologists are in the process of creating registries for names, nomenclatural acts, and related information (Pyle and Michel 2008). New submissions could be vetted, with logic reasoners configured to enact nomenclatural rules, resulting in improved quality control (Patterson et al. 2016). (4) Lastly, due to the complex interactions between evolving taxonomic perspectives and nomenclatural emendations, the ways in which such changes are implemented can have varying effects on nomenclatural stability (e.g., van der Linde and Houle 2008; ICZN 2010; Nicholson et al. 2012). Whenever alternatives exist to reconcile taxonomic and nomenclatural changes, logic reasoners can be deployed to maximize nomenclatural stability or other criteria (Vences et al. 2013). In summary, representing nomenclatural rules in computational logic may yield both theoretical and practical benefits.

What do we mean by “nomenclatural rules”? To clarify, the domain-specific Codes of nomenclature are not a monolithic body of rules applicable to all life (Hawksworth 2001; David et al. 2012). The particularities of each rule set, combined with the histories of certain taxonomic names, can lead to unique complexity challenges, requiring expert knowledge to attain nomenclatural resolution (e.g., Vane-Wright 2003; Rhodin and Carr 2009). However, in spite of inter-Code differences and sometimes bewildering use cases, there are several overarching principles of Linnaean nomenclature – such as Priority and Typification – that can be translated into general, conditional statements and thus become tractable for logic-based reasoning. our present focus is on representing several of these foundational nomenclatural principles in logic.

Here we create a hypothetical 20 nomenclatural taxon use case and logically model its transition in following a specific taxonomic action. The starting conditions embody several core principles (henceforth capitalized) of the International Code of Zoological Nomenclature (ICZN 1999). These include: Binominal Nomenclature (Article 5), Priority (Article 23), Coordination (Article 36), Homonymy (Article 52), Typification (Article 61), and the structural requirement of Gender Agreement (Article 31.2). These starting conditions are encoded into an Answer Set Programming (ASP) language and program (Gelfond and Lifschitz 1988; Gelfond 2008; Brewka et al. 2011). In a subsequent step, modeled with a transition system approach (Lifschitz and Turner 1999), one of 12 species-level names (child) and the corresponding type specimen are transferred to another genus name (parent). Permutations of this child-to-parent reassignment action lead to 36 alternative scenarios, where each ending condition requires 2-14 logically contingent nomenclatural emendations (Tables 1 and 2). We show that an ASP reasoner (Gebser et al. 2011) can correctly infer the Code-mandated changes for each input and transition scenario, and visually output the ending conditions. In the Discussion, we assess the outcomes in relation to a broader discourse about the interaction of nomenclature, taxonomy, and logic-based representations of systematic change.

## Materials and Methods

To our knowledge this is the first application of an ASP transition system approach to model an inference process in the nomenclatural-taxonomic domain (though see Brooks et al. 2007; Gebser et al. 2008; and Franz et al. 2016a; for different uses of ASP in biology). We regard ASP as an alternative or complementary solution to the challenge of knowledge representation and reasoning in biology, where Description Logics is the prevailing paradigm (Grenon et al. 2004; Smith et al. 2007; Baader et al. 2008). To bring the readership into this field, we first describe the properties of ASP in relation to the inference needs for the use case. This review is sets the stage for modeling of the nomenclatural change scenario.

### Properties of the Answer Set Programming Approach

The term Answer Set Programming was coined in the late 1990s for a declarative form of logic programming rooted in non-monotonic reasoning and stable model semantics (Lifschitz 2008; Eiter et al. 2009; Brewka et al. 2011). Problems are solved in ASP by specifying sets of rules and constraints and then generating stable models that satisfy these. The stable models are inferred by grounding the specified input domains, variables, and conditions during the reasoning process. The latter generates all answer sets that are not logically prohibited by the joint constraints (Fig. 2).

The ASP approach has several desirable properties in connection to the complex rules of nomenclature, as follows. (1) Answer Set Programming uses an expressive language resembling that of other Semantic Web languages (Eiter et al. 2008). (2) There are powerful ASP reasoners (answer set solvers) such as the Potassco Answer Set Solving Collection (Gebser et al. 2011). (3) Unlike Description Logic, the approach is based on the closed world assumption, which permits deciding that conditions not proven to be true are false (“negation as failure”; Gelfond and Lifschitz 1988). (4) It offers elaboration tolerance, i.e., the ability to retain reasoning abilities while taking into account additional constraints (McCarthy 1998). (5) It also supports default reasoning, i.e., the ability to stipulate the truth of certain conditions unless specified otherwise (Reiter 1980). (6) Answer Set Programming can represent transition systems through automated reasoning about action domains, by generating plans for such domains. (7) The approach is translatable to other logic paradigms such as First-Order Logic and Description Logic (Eiter et al. 2008; Kutz et al. 2010). Jointly these properties make ASP highly suitable for performing complex, exception-rich, rule-based reasoning in the biological domain. However, (8) at present most ASP applications lack an end user interface as mature as (e.g.) the Protege platform (protege.stanford.edu/), instead requiring knowledge of command line and logic program coding.

### Starting Conditions

To set the stage for this use case, we must first specify the nomenclatural and taxonomic boundaries and reasoning intentions (Figs. 1 and 2; Tables 1 and 2). The ICZN (1999) contains 90 articles. Of these, only a subset can be represented here. We also recognize that certain nomenclatural changes are triggered independently of new taxonomic insights. Conversely, not all taxonomic changes are mirrored in nomenclatural adjustments (Franz 2005; Franz et al. 2008;, 2016a, 2016b Remsen 2016). The various dependencies between nomenclatural and taxonomic emendations make it necessary to characterize the *taxonomic* starting and change conditions for the use case as unambiguously as possible. We thereby model specific principles and articles of the IZCN that guide the reasoning process towards succinct ending conditions.

**Fig. 1.**
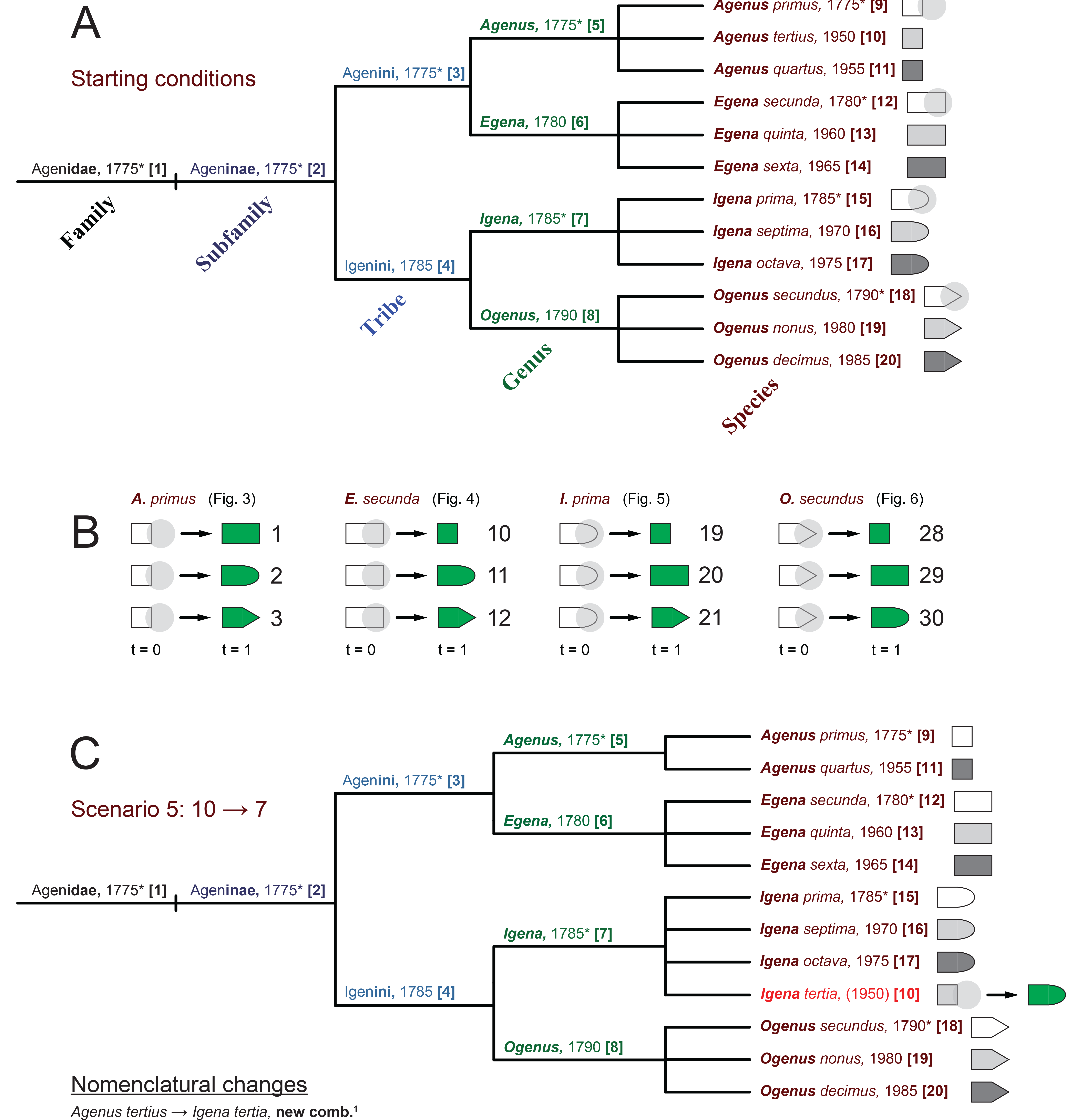

**Fig. 2.**
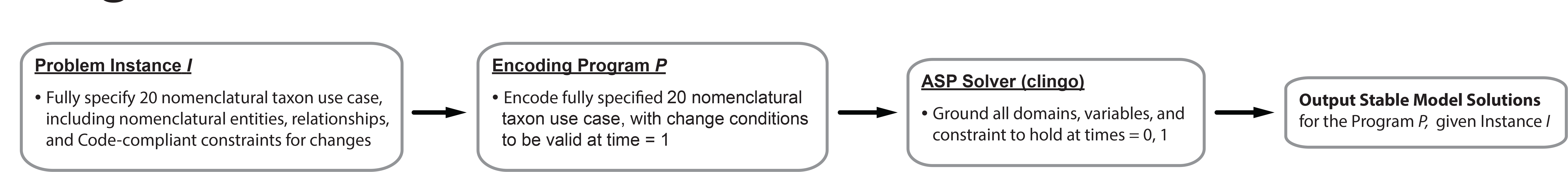

The taxonomic names themselves are another set of variables needing prior specification. Given a taxonomic change scenario, the reasoner can infer *that* certain nomenclatural emendations are necessary. However, the reasoner has no capacity to conceive *which* name strings should be used. For instance, under the proper rules and change conditions, the program can infer that the species-level epithet *tertius* will require replacement with a new, valid, masculine genus name. But it cannot decide whether the name string *Zyzzyzus* (Brinckmann-Voss and Calder 2013) is preferentially suited for this purpose in terms of availability, Latin grammar (Casadio and Lambek 2005), and perhaps also cognitive or aesthetic considerations. Consequently, we must provide the reasoner upfront with a set of Code-compliant name options that can be applied – as logically required – to conclude the transition.

The use case entails two temporal conditions or steps (t = 0, 1). One condition occurs prior to the taxonomic change (t = 0), and the other condition occurs subsequent to the taxonomic change (t = 1). The pre-change phase is set in the year 1985, whereas the post-change phase is set in the year 2000. Prior to the change, 20 ranked, nomenclaturally connected names exist: one each at the family and subfamily level (endings *-idae* and *-inae*); two at the tribal level (-*ini*); four at the genus level; and 12 at the species level (Fig. 1A). Each name is assigned a unique numerical identifier, displayed in square brackets: [1], [2],…[20]. Lower-level child names are allocated in equal numbers among their respective parent names, i.e., each generic name contains three species names, and each tribal name contains two generic names. The 12 species-level names were validly published in two preceding time clusters, referred to as (1) the “type period” – extending from 1775 to 1790 – and (2) the “non-type period” – extending from 1950 to 1985. The publication years for species-level epithets are specified such that each generic name accommodates exactly one epithet from the earlier type period and two epithets from the later
non-type period. No two species-level names were published in the same year, thereby rendering judgments of Priority unambiguous.

As a consequence of Coordination, the distribution of type period publication years at the species level generates four distinct lineages of Priority; viz. (1) *Agenus primus*, 1775 – [9], [5], [3], [2], and [1]; (2) *Egena secunda*, 1780 – [12] and [6]; (3) *Igena prima*, 1785 – [15], [7], and [4]; and *Ogenus secundus*, 1790 – [18] and [8]. Two generic names are masculine ([5], [8]) and two are feminine ([6], [7]), respectively. Moreover, two pairs of epithets share the same Latin root, either *prim-* ([9], [15]) or *secund-* ([12], [18]). This circumstance has implications for Homonymy under certain change scenarios.

Each of the species-level names is anchored by a single, vouchered, unambiguously assigned type specimen (Fig. 1A). Note, however, that “assignment” is used here in the nomenclatural sense, i.e., there is agreement which specimen is being referred to (Witteveen 2015, 2016). Such an unambiguous nomenclatural assignment differs from agreeing on the taxonomic identity, or more precisely, the interpretation of certain taxonomic traits that the nomenclatural type specimen in question may or may not exhibit according to an expert’s re-/assessment.

Because the logic of reasoning ultimately falls back on the perceived taxonomic identity of type specimens, we need to stipulate traits for the latter as well (Fig. 1A). Accordingly, each type specimen within the same genus-group name varies in one species-level taxonomic trait related to its pigmentation: (1) unpigmented, (2) lightly pigmented, and (3) strongly pigmented. In addition, all three type specimens assigned to the same genus-group name *prior* to the change share a second, genus-level diagnostic feature, as follows: (i) the type specimens for the three *Agenus* species names have a square shape; (ii) those for the three *Egena* species names have a rectangular shape; (iii) those for the three *Igena* species names have a hemi-elliptical shape; and (iv) those for the three *Ogenus* species names have a pentagonal shape. The combination of three species-level and four genus-level diagnostic traits establishes each of the 12 type specimens as taxonomically unique in the original assessment. Jointly the above conditions specify the pre-change status of the use case, which is complete as of 1985.

### The Change – Type Specimen Reinterpretation

The transition from the pre- to the post-change phase is triggered by the *reinterpretation* of the taxonomic identity of one individual type specimen (Fig. 1B). This reinterpretation occurs in the year of 2000. Recall that the respective shapes of the four specimens anchoring the type period epithets (1775-1790) – square, rectangular, hemi-elliptical, and pentagonal – formed the diagnostic foundation for delimiting the corresponding genus-level taxon (concept) to which additional species-level entities were subsequently assigned. These additions were performed under the assumption of taxonomic correspondence, thus effectively reconfirming the original shape descriptions. However, in 2000 matters are reassessed with a different outcome. The original interpretation of a type specimen’s diagnostic feature is now found to have been erroneous, creating a case of “reference by misdescription” (Donnellan 1966). Instead, the type specimen is diagnosed to exhibit one of the *other* shapes present in species-level entities assigned to one of the respective three generic entities (Fig. 1B).

While the example is contrived, in practice such type reinterpretations are not infrequent (e.g., Ribot et al. 1996; Woodley et al. 2011; Cappellini et al. 2013; Laloy et al. 2013; Witteveen 2015). Reassessments of the taxonomic identity of type specimens are very frequently involved in determining heterotypic synonymy. In the present use case, the type reinterpretation requires an adjustment that achieves taxonomic congruence across the named entities such that similarly shaped type specimens are assigned to the same genus-level name.

The reinterpretation of the type specimen has specific consequences. In particular, the 1:1 cardinality relationship between type specimens and valid species-level epithets is *not* altered through the transition, remaining at 12:12 for each phase. With regards to taxonomic or topological congruence, a genus that previously contained three species now entails one species less, whereas another genus entails one additional species. Considering the 3-3-3-3 numerical allocation of specific to generic entities in the pre-change phase (Fig. 1A), this means that exactly two of these numbers shift to 2 and 4, respectively. Because the asserted shape of the type specimen anchoring each of the 12 species-level epithets is potentially ‘corrigible’ to match any of the three alternative shapes (Fig. 1B), there are 36 possible ending conditions for the use case (12 type specimens × 3 alternative shapes).

### Ending Conditions

Each of the 36 ending scenarios produced in the year of 2000 (t = 1) requires application of a unique set of logically induced, Code-mandated nomenclatural rules, and therefore triggers specific but different nomenclatural emendations. The particular rules and changes to be enacted depend on the individual type specimen under reinterpretation and the morphological shape newly ascribed to this specimen. Differential sets of principles and articles in the ICZN (1999) are invoked accordingly.

The outcomes fall into two broad categories. Twenty-four of the 36 ending conditions are caused by reinterpretations of type specimens anchoring eight species-level epithets that were created in the later (non-type) phase; i.e., [10], [11], [13], [14], [16], [17], [19], and [20]. These names *lack* Priority in relation to other same-ranked names typifying *both* the source and target genus-level names between which the names transition. Thus, each of these transfers requires only a limited set of nomenclatural changes; specifically, the creation of one new combination (Fig. 1C).

The remaining 12 transition scenarios are nomenclaturally more complex (Figs. 3–6). Here the act of reinterpretation affects type specimens anchoring one of four species-level epithets created in the earlier (type) phase; i.e., [9], [12], [15], and [18]. Priority considerations now *drive* the nomenclatural outcomes through multi-level dependencies. First, the epithets whose types are reinterpreted have Priority over each of the two epithets accommodated under the respective (starting phase) genus-level name (Fig. 1A). We may call this kind of Priority relative to initially congeneric epithets “local Priority”. An example of an epithet with local Priority is *secundus*, 1790 – relative to *nonus*, 1980, and *decimus*, 1985. “Global Priority”, in turn, holds throughout the entire nomenclatural hierarchy. The epithet *primus*, 1775, holds this status among all 12 epithets. Hence, in deciding on the appropriate nomenclatural emendations for the 12 more complex cases, one must assess the relative Priority among paired epithets that each have local Priority; e.g., (1) *primus*, 1775, versus (2) *secunda*, 1780; or (1) *prima*, 1785, versus (2) *secundus*, 1790, etc. Whichever epithet lacks Priority in the pairwise comparison effectively transitions into a new combination, where the post-transition genus-level name is that corresponding to the Priority-carrying epithet.

**Fig. 3.**
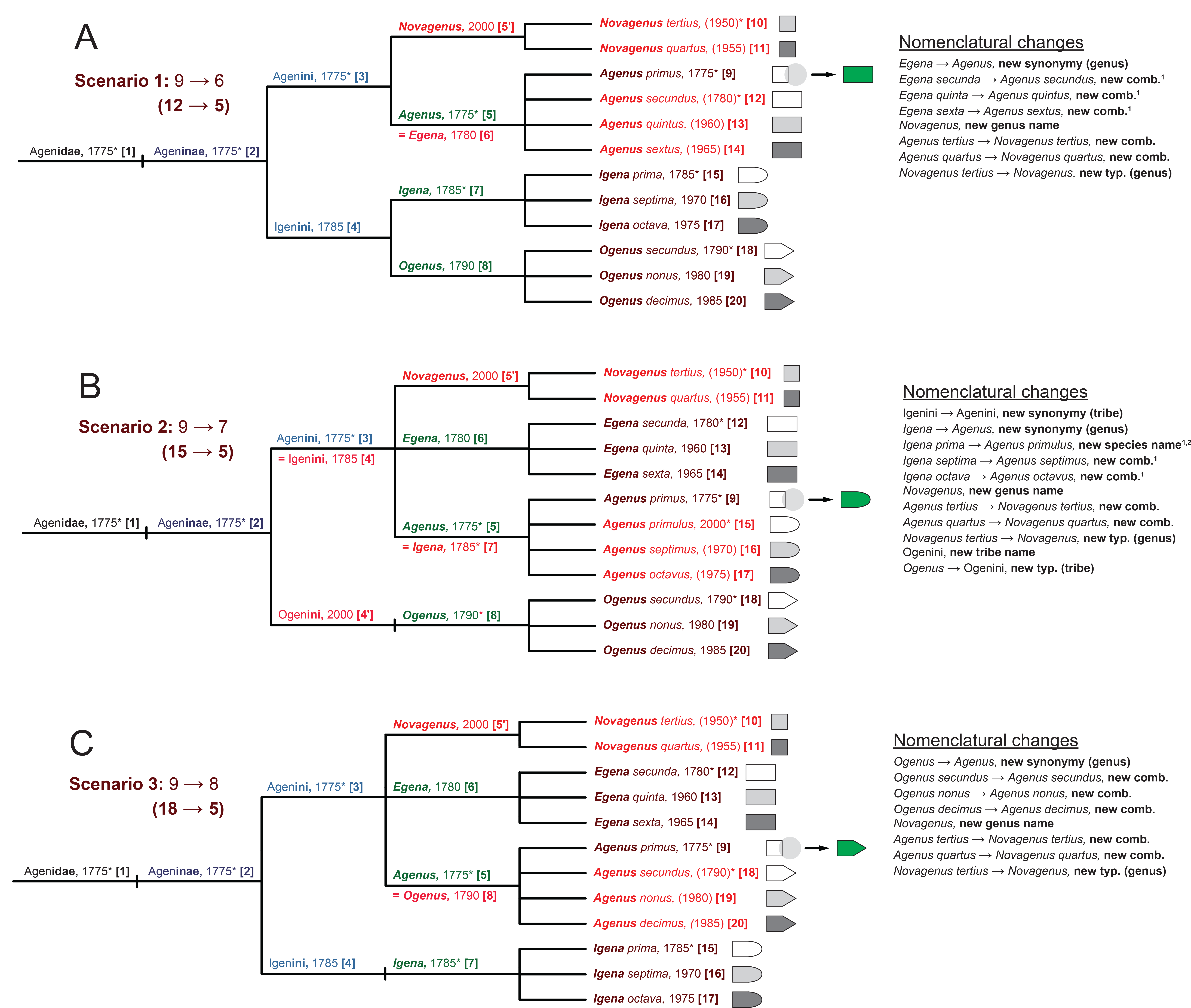

Second, we need to account for the principle of Coordination, which further complicates matters. Consider an epithet with initial, local Priority – e.g., *secundus*, 1790 – in relation to another such epithet, which has Priority in the more expansive context – e.g., *secunda*, 1780 (Fig. 4C). In such cases the two epithets’ respective genus-level names – *Egena*, 1780, and *Ogenus*, 1790 – also enter into a synonymy relationship: only one name remains valid. In some instances, the coordinated tribe-level names must enter into synonymy as well; see ICZN (1999), Article 40.1. In addition, certain epithets that originated in the non-type period will newly acquire local Priority, and thereby trigger the creation of new genus- and/or tribe-level names, new typifications of these, and new genus name/epithet combinations (e.g., Fig. 4C).

**Fig. 4.**
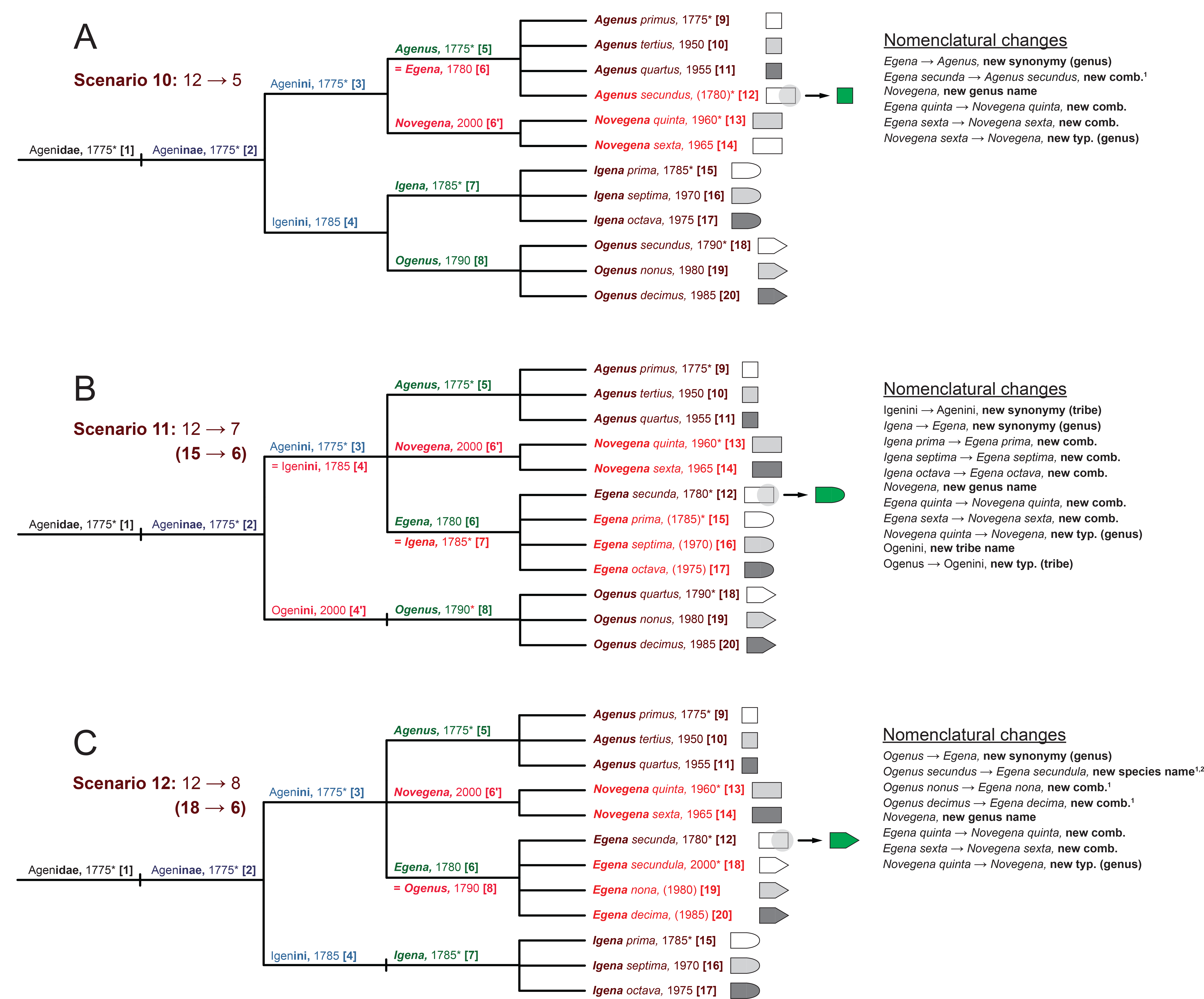

Lastly, no taxonomic change in the use case can affect the validity of the nomenclatural lineage of Agenini, 1775, to *Agenus primus*, 1775 ([3] → [5] → [9]), which has global Priority (Fig. 3). In the 3/12 complex cases where the type specimen anchoring the epithet *primus*, 1775, is reinterpreted to have a non-square shape, this novel insight will *not* affect the validity of the globally Priority-carrying name lineage. Instead, it is required that the other lineage in the comparison, having local Priority, move into synonymy. The two additional, non-type period epithets subsumed under that latter lineage are thus also newly combined with the genus-level name *Agenus.* Moreover, the two names *Agenus tertius*, 1950, and *Agenus quartus*, 1955, no longer represent valid combinations (given their square-shaped types), and trigger the creation and typification of a new genus-level name. Thus, even though the name *Agenus primus*, 1775, is nomenclaturally stable across all 36 scenarios, reinterpretations of the type specimen anchoring this name can have considerable and cascading nomenclatural effects for other names and relationships modeled in the use case (Fig. 3).

### Documentation of Logic Approach and Outcomes

We assume that the ASP approach is novel to most readers, and therefore make an effort to present the methods and results in an accessible way. Our analysis entails two main components: (1) the specification of a transition system with starting and ending conditions that require application of multiple principles of zoological nomenclature (Figs. 1 and 2); and (2) the translation of these conditions and action rules into a logic program that an ASP reasoner can analyze to generate the 36 possible outcomes (Tables 1 and 2). The finer details of our logic program are likely not of great interest to the systematics readership, and are therefore presented in the Supplementary Materials S1-S5.

The code for this program is written in the language F2LP (Lee and Palla 2009), and comprises less than 600 lines including annotations (Supplementary Materials S1). F2LP is a logic programming language that permits encoding sets of First-Order Logic formulas into stable model semantics suited for an ASP reasoner. Our program is written to interface with software tools produced by Potassco – the Potsdam Answer Set Solving Collection (Gebser et al. 2011). In particular, the code is processed with the system *clingo*, which combines the variable grounder *gringo* with the Answer Set solver *claspD* to process disjunctive logic programs (Gebser et al.2015).

The output inferred by the system contains both hierarchical taxonomic and nomenclatural information; in particular: (1) relationships (edges) among nodes in the single starting tree and set of ending trees, and (2) information on the names and nomenclatural relationships, actions, and changes that reflect the ending conditions at each node. To visually represent and confirm the correctness of the outcomes, we configured a simple Java program that translates the *clingo* text string output into .dot files. The latter can be displayed as taxonomic trees with edges and named nodes in the open source GraphViz visualization software (Gansner and North 2000). The nomenclatural (relationship) information is output directly in textual format by *clingo*.Information on the corresponding *clingo* commands, textual reasoner output, Java visualization program, and generated GraphViz visualizations is provided in the Supplementary Materials S2-S5, respectively – all are available via the Dryad Digital Repository.

The Results are separated into two main sections. The first of these focuses on presenting the principal components and actions of the logic program in a high-level pseudocode. This nontechnical code review explains how our ASP approach succeeds in representing the taxonomic and nomenclatural conditions and transitions of the use case. We thereby illustrate to the readership how specific nomenclatural principles such as Priority or Coordination are encoded in ASP logic. The second section presents the use case outcomes, emphasizing the relationship of nomenclatural and taxonomic changes across the 36 transition scenarios. To this end, we aggregate the numbers of new combinations, new names (species, genus, tribe), new synonymies (genus, tribe), and new typifications (genus, tribe), at the respective taxonomic ranks. For new combinations and new species names, we also report on the number of cases where application of Gender Agreement requires changing the ending of an epithet from masculine to feminine, or vice-versa.

While the present analysis is merely an exploratory step, the compiled numbers provide some general insights into the logical interdependencies of nomenclatural starting and ending conditions in light of a change in taxonomic assessment. As outlined in the Discussion, a better understanding of these dependencies may eventually lead to logic-enabled optimizations of nomenclatural and taxonomic change actions in biodiversity data environments.

## Results

### Documentation of the Answer Set Programming Approach

The general structure of our ASP programming approach is provided in Table 1. All sections are necessary to run the code with *clingo* and obtain the desired outcomes. However, because the ASP reasoner evaluates all constraints simultaneously, the sequence of the sections and subsections is relevant only to enhancing human readability of the code. Accordingly, Section 1 defines the overarching transition system domains, variables for subsequent if-then constraints, and starting conditions; including two steps, 20 nodes, all node-associated ranks, node-associated names, Priority-constrained name lineages, mono- and binomial names, and name-associated years of publication. An important feature allowing further elaboration of this code is the semi-independent modeling of (1) name identities and relationships (’nomenclature’), and (2) node identities and relationships (’taxonomy’); where “semi-independent” corresponds to the complex cardinality interactions that occur in nomenclatural/taxonomic change scenarios (e.g. Vane-Wright 2003; Franz 2005; Remsen 2016).

**Table 1:** High-level ASP pseudocode for the 20 nomenclatural taxon use case. The actual F2LP code, input commands, and output results (clingo text, GraphViz visualizations) are presented in the Supplementary Materials S1-S5, respectively.

| Section | Constraint or action modeled in Answer Set Programming logic |
| --- | --- |
| **1.** | **Define the transition system domains, variables, and starting conditions** |
| 1.1. | Define a system that has two phases or steps: 0 = original; 1 = revised. |
| 1.2. | Define that the system has 20 nodes, labeled node 1 to node 20. One or more nodes can share the same rank (family to species). |
| 1.3. | Define that rank-specific nodes are part of the entire set of nodes. |
| 1.4. | Specify that higher-level names are constrained by coordinated (type-lineage) identities (*Coordination*). Generic names and epithets are constrained by gender identities and corresponding spellings (*Gender Agreement*). |
| 1.5. | Define multiple domains for nodes: generally, and for each taxonomic rank (family to species). |
| 1.6. | Specify the original tree shape at time = 0, with 19 edges connecting the 20 nodes. |
| 1.7. | Specify the original generic name/epithet combinations for the corresponding nodes at time = 0, with the corresponding publication years and gender identities. |
| 1.8. | Constrain each epithet to remain assigned to its original node; thereby, species-level node numbers remain consistent throughout the two phases of the transition scenario. |
| 1.9. | Specify the coordinated (type-lineage) identities for sets of names – including each name's prefix and rank-specific endings – for the starting and ending conditions. |
| 1.10. | Constrain the publication year for each name, including new names created in the year of 2000 (though excluding epithets which were previously specified in 1.7.) |
| **2.** | **Specify the globally applicable uniqueness and existence constraints** |
| 2.1. | Constrain that each node is assigned a single, rule-compliant name – either mono- or binomial – as well as the year and the edge to the parent node, excepting the root node which has no parent node. |
| **3.** | **Specify the globally applicable inertia conditions for entities and relationships that remain consistent *unless* change is mandated by the transition model** |
| 3.1. | Constrain that all edges, node-name-year-gender assignments, and node years specified as valid at time = 0 will remain so at time = 1, ranging from the family level down to the genus level (however, species-level edges may change). |
| **4.** | **Specify that the Priority of an epithet-level node and its coordinated higher-level nodes is established by having the more senior publication year in comparison to another epithet-level node (*Priority*)** |
| **5.** | **Assign nodes persistently to their respective names and years at each taxonomic rank (family to species)** |
| **6.** | **Infer the rule-mandated nomenclatural changes at time = 1, for persisting and newly valid names (ending conditions)** |
| 6.1. | Constrain that the valid generic name (at any time) has the senior, Priority-carrying epithet. |
| 6.2. | Constrain that epithets persist from time = 0 to time = 1; however, the corresponding generic name is determined by the identity of the senior, Priority-carrying epithet. |
| 6.3. | Infer that an epithet is combined with a new generic name *if* it newly acquires Priority at time = 1 through the transfer of a previously Priority-carrying epithet (at time = 0). |
| 6.4. | Exception to 6.3.: The node of the more senior, Priority-carrying epithet is *not* transferred in the transition from time = 0 to time = 1. Instead, another, not Priority-carrying node is transferred "inversely". |
| 6.5. | Infer that the tribe-level name is coordinated (newly, if required) with the Priority-carrying genus-level name. |
| 6.6. | Infer that the subfamily-level name is coordinated (newly, if required) with the Priority-carrying tribe-level name. |
| 6.7. | Infer that the family-level name is coordinated (newly, if required) with the Priority-carrying subfamily-level name. |
| **7.** | **Specify the available new names and their respected years of publication, as required by previous constraints** |
| **8.** | **Specify the changed taxonomic conditions (stable models) that are valid at time = 1: choose exactly one species-level node to be transferred as the child of another parent genus-level node than that which it had at time = 0** |
| 8.1. | Select one species-level node to change its parent genus-level node at time = 1. |
| 8.2. | Exception to 8.1.: The change is not executed if the new node placement affects the same original edge (i.e., no effective change). |
| 8.3. | Establish that the new placement of the species-level node *creates* an edge to the corresponding genus-level node. |
| **9.** | **9. Identify and aggregate the required nomenclatural changes at time = 1** |
| 9.1. | Record a *new combination* when the node names at times = 0 and 1 are not identical. |
| 9.2. | Record a *new placement* strictly in recognition of the relative Priority of locally and globally senior children; in particular, the latter are *not* transferred (as specified in 6.4.). |

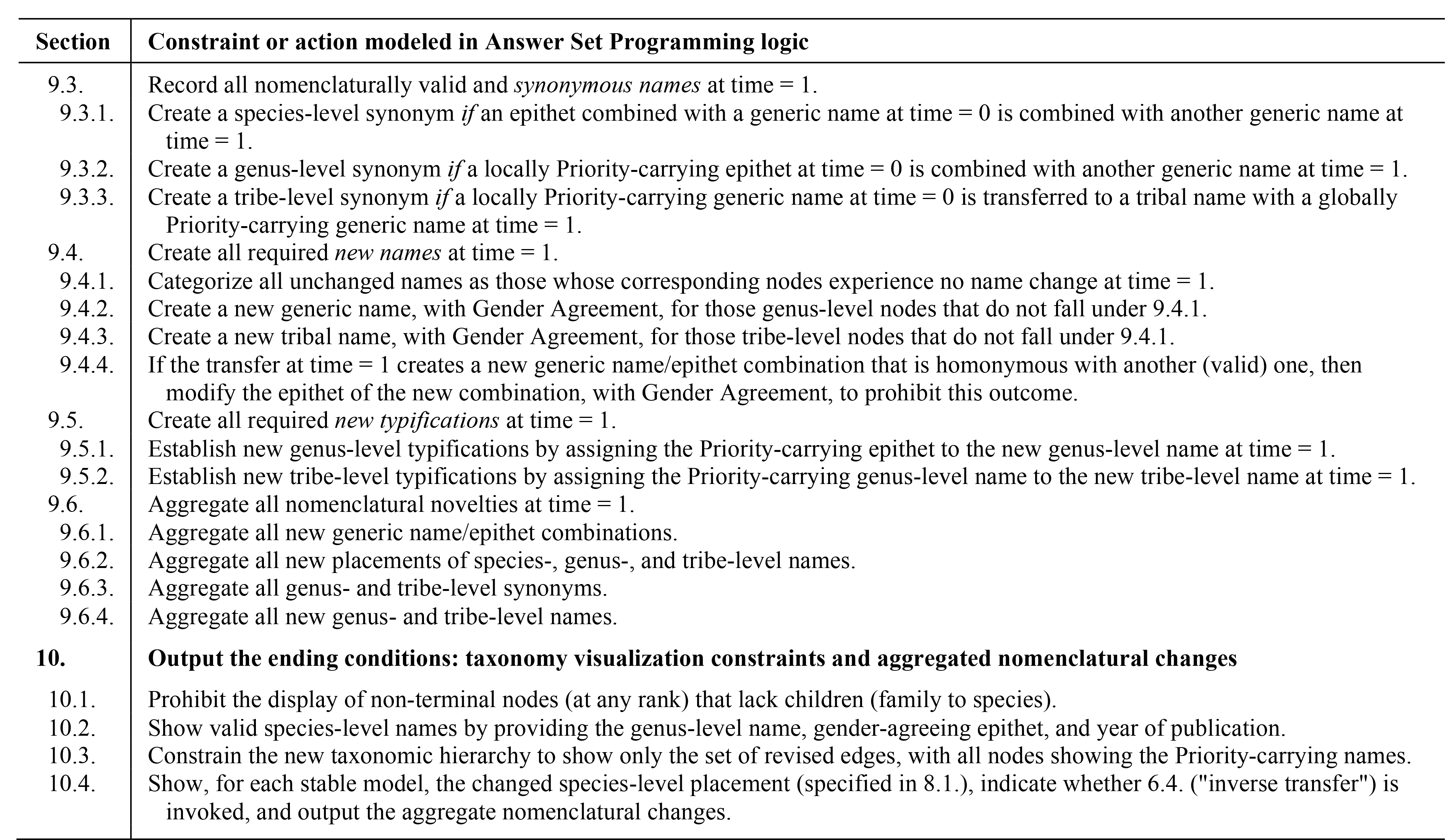

Sections 2–5 establish several globally applicable conditions, i.e., conditions that must hold true for any step in the transition system. These include: assignment of a single valid name to each node, with rank, year, and parent, excepting the root node; inertia conditions – all t = 1 conditions are equal to the t = 0 conditions *unless* the change triggered by the species-level node move overwrites this constraint; Priority among children nodes that have the same parent node; and persistence of taxonomic ranks throughout the transition.

Sections 6-8 represent the key rules needed to compliantly model the taxonomic change and nomenclatural emendations to be enacted at t = 1. The core change itself is specified in Section 8.1., by simply requiring one species-level node to not have the same parent node at t = 1 which it had at t = 0. This constraint in effect creates a new species-to genus-node edge at t = 1, while keeping the total number of nodes consistent. Sections 6 and 7 establish which existing and new names are held as valid at t = 1. These rules specify how the Priority (and hence validity) of a genus-level name is determined by the most senior epithet associated with that generic name at time = 0. They furthermore allow epithets to ‘newly become the most senior’ for the respective parent genus name at t = 1, if the taxonomic change (see above) triggers this transition. However, this latter constraint – and others logically tied to it – has an important exception (Section 6.4.): if the mandated node move *violates* the relative or global Priority status of genus names and species epithets as established at t = 0, then an ‘inverse move’ is executed (e.g., Fig. 3). This means that, in spite of the reassignment of a child node to a new parent node, the globally Priority-carrying binomial – e.g., *Agenus primus*, 1775, in all Fig. 3 scenarios – is maintained as valid. Moreover, the other locally Priority-carrying binomial will ‘move into *Agenus'*, even though neither its type specimen nor those of other congeners recognized at time = 0 were reinterpreted to exhibit another shape (hence we say “inverse”, as in: initially intended to affect the status of another node/name association). The remainders of Sections 6 and 7 constrain that Coordination propagates from the lowest to the highest represented taxonomic rank, and that any new name(s) requiring creation at t = 1 should start with the letters *Nov-* prefixing the previously valid name. For instance (Fig. 5), *Novigena*, 2000, is used as a new genus name for valid epithets (*septima, octava*) previously associated (t = 0) with the now synonymized genus name *Igena* (t = 1). Lastly, if the move of a node and its associated epithet is bound to create two (binomial) homonyms, then this Code violation will be *prohibited* by adding the endings *-ulus* (e.g., *primulus* instead of*prim–us)* or *–ula* (e.g., instead of *secund-a)* to the epithet that will no longer have local Priority at t = 1 (see, e.g., Figs. 4C and 5A). These cases are not recorded as new combinations, but instead are new replacement names, and are recognized as new species-level names in Table 2.

**Fig. 5.**
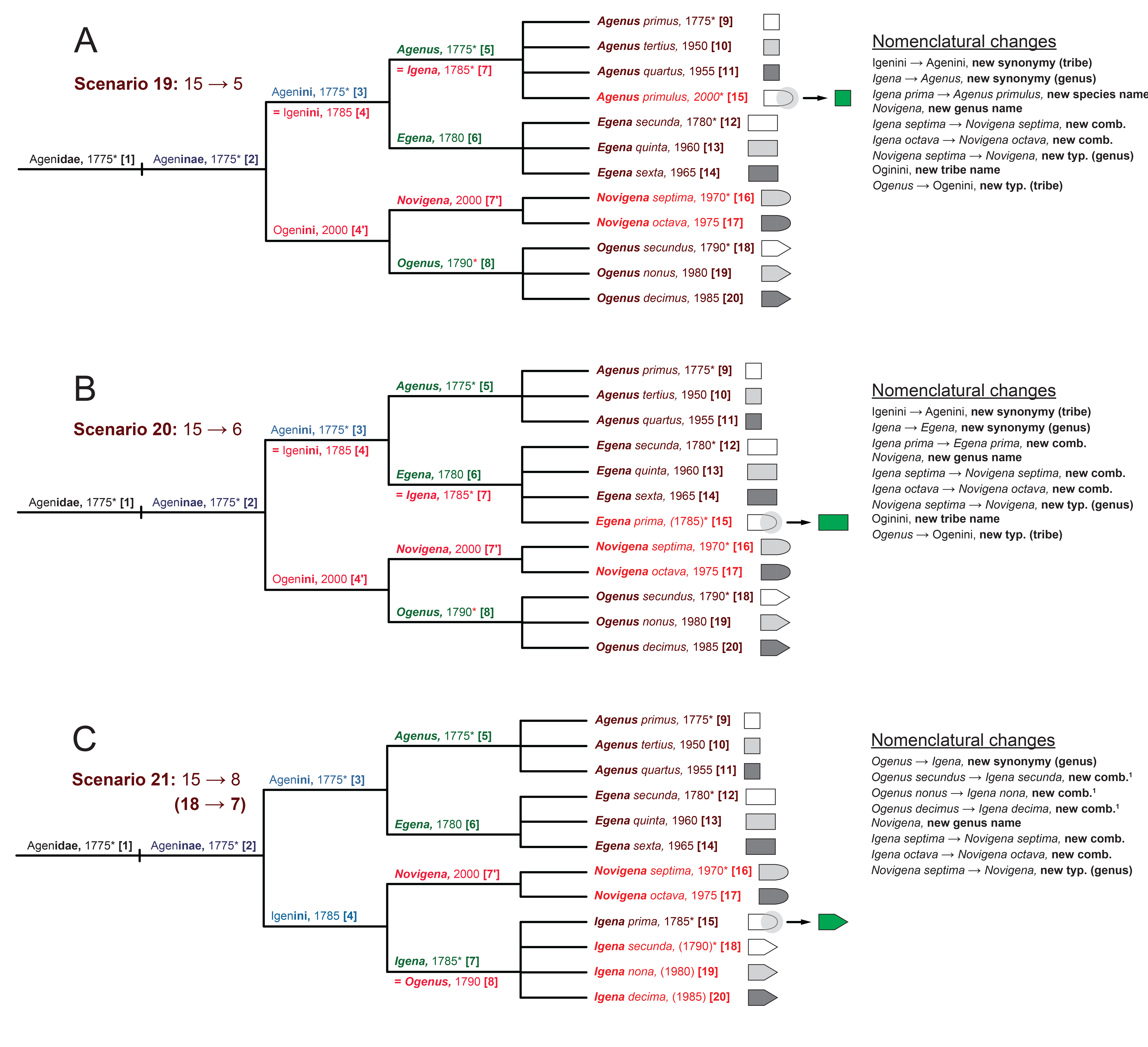

Sections 9 and 10 respectively identify all resulting nomenclatural emendations and generate the corresponding output topologies, valid nodes names, and nomenclatural status and relationship updates. The code of Section 9 includes if-then constraints that identify at t = 1: new combinations and concomitant applications of Gender Agreement leading to new epithet endings; new synonymies at the generic and tribal levels; new species-, genus-, and tribe-level names; and new typifications of higher-level names by their entailed, newly senior lower-level names. The Section concludes with commands that aggregate totals for each of these emendations for the corresponding stable model. The final Section 10 provides code to output each possible node topology as a set of “revised edges”, the valid node names for each rank, as well as the list of nomenclatural emendations. If an inverse move is indicated in the model, this information is also printed out (see Supplementary Materials S5).

### Analysis of Nomenclatural Transition Scenarios

The ASP programming approach correctly generates all 36 transition scenarios (stable models) and the required nomenclatural changes (Figs. 1C, 3-6; Table 2). Given the use case constraints, exactly six of the 36 scenarios require an inverse move: three involving the node/binomial *Agenus primus*, 1775 (Figs. 3A-3C); two concerning the node/binomial *Egena secunda, 1780* (Figs. 4B and 4C); and one including the node/binomial *Igena prima*, 1785 (Fig. 5C). The node/binomial *Ogenus secundus*, 1790, does not have Priority over the three aforementioned names, and thus cannot trigger an inverse move (Fig. 6).

**Fig. 6.**
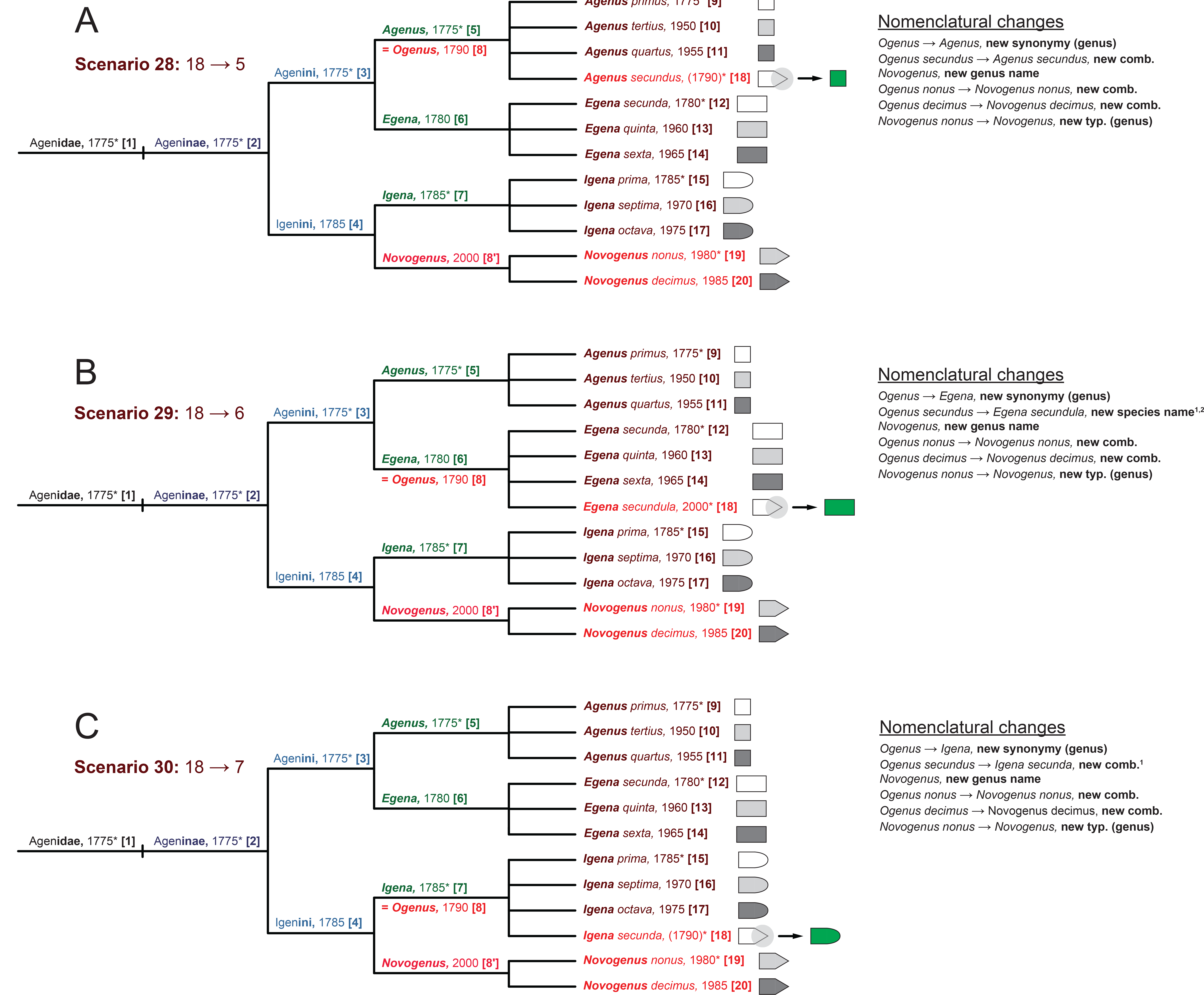

The 12 complex scenarios involving reinterpretations of specimens designated during the type period (1775-1790) account for 114/154 (74%) of the recorded nomenclatural emendations at t = 1 (14 of these are changes of epithet endings to achieve Gender Agreement). This corresponds to an average of 9.5 changes per scenario. As many as five new combinations and/or new species-level names are triggered in several of these scenarios. In contrast, the 24 remaining scenarios involving reassessments of non-type period specimens require only 40/154 changes in nomenclature (26%) – all singular new combinations (16 of which also require epithet changes to comply with Gender Agreement) – for an average of 1.7 changes per scenario.

The nomenclatural particularities of the 12 complex scenarios are well documented in Figs. 3-6 and require little additional explanation. Each scenario requires the creation and typification of a new generic name at t = 1, as well as a genus-level synonymy (Table 2). In four cases, the node transfer affects an epithet associated with a source genus-level name (either *Igena prima*, 1785, or *Ogenus secundus*, 1790) sat t = 0 that is homonymous with that of another epithet associated with the target genus-level name *(Agenus primus*, 1775, and *Egena secunda*, 1780, respectively). These cases require the creation of new species-level replacement names in accordance with Priority, resulting in the new epithets *primulus*, 2000 (Figs. 3B and 5A) or *secundula*, 2000 (Figs. 4C and 6B). In two of these scenarios (Figs. 3B and 5A) and two others (Figs. 4B and 5B), *Igena*, 1785, is synonymized at t = 1 with another generic name in the Agenini, 1775 (either *Agenus* or *Egena*). Consequently, and in compliance with Coordination, the tribal name Igenini, 1785, is also synonymized with Agenini. This means that a new tribal name Ogenini, 2000, must be created and typified with the generic name *Ogenus*, 1790. These scenarios have the most wide-ranging consequences in terms of requiring the application of diverse set of nomenclatural principles and rule.

### Complementary Representation Approach – Taxonomic Concept Alignments

Elsewhere we have applied ASP to promote the taxonomic concept approach (Franz et al. 2005, 2016a, 2016b). This approach represents each published taxonomy (t = 0, 1) separately, via *taxonomic concept labels* (name sec. author; where sec. means “according to“) and parent-child *(is_a)* relations, and then employs expert-provided Region Connection Calculus (RCC-5) *articulations* to express taxonomic congruence or non-congruence (inclusion, overlap, exclusion) between regions pertaining to multiple taxonomies. An ASP reasoning toolkit (Chen et al. 2014) is available to (1) check for the logical consistency of these input constraints under certain taxonomic covering assumptions, (2) infer additional implied articulations, and (3) visualize consistent multi-taxonomy *alignments.* The latter may be viewed as taxonomic *meaning resolution* maps that can guide information integration across the varying taxonomies. In this sense, the alignments can complement representations of ‘identity in light of change’ as provided by Linnaean names and nomenclatural relationships (homonymy, synonymy). Simple RCC-5 metrics are also suitable for assessing the performance of taxonomic names as identifiers of congruent or non-congruent taxonomic meanings (Franz et al. 2008, 2016a, 2016b).

Because the transition scenarios of this use case are strongly and heterogeneously affected by the application of Priority, Coordination, Homonymy, Typification, and Gender Agreement – all key principles in the Linnaean tradition – it is illustrative to model the starting and ending conditions as RCC-5 alignments of taxonomic concept hierarchies. The complete set of 36 alignments, each with relevant input and output data files, is provided in the Supplementary Materials S6-S8.

Four exemplary alignments are shown in Figs. 7 and 8, corresponding to scenarios 1, 2, 4, and 5. Table 3 summarizes the alignment patterns obtained for the 36 scenarios. Interestingly, there are only three general patterns that can be categorized as follows: either a species-level node is transferred among genus-level nodes placed under two tribe-level nodes at t = 0 (Agenini *and* Igenini, respectively), and in that case it is either (1) an intended move or (2) an inverse move; or (3) the transfer involves genus-level nodes placed within in the same tribe-level node at t = 0 (Agenini *or* Igenini). Pattern (1) is exemplified by scenario 5 and shown in Fig. 7A (compare with Fig. 1C). This pattern holds for 20 scenarios, and in each case corresponds to alignments that have 15 congruent regions – 3 above the species level – and 6 overlapping articulations. Pattern (2) is true for scenario 2 and shown in Fig. 7B (compare with Fig. 3B). This pattern is present in 4 scenarios, where the alignments have 15 congruent regions and 4 overlapping articulations. Lastly, pattern (3) is represented by scenarios 4 and 1 and displayed in Figs. 8A and 8B (compare the latter with Fig. 3A). The pattern applies to 12 scenarios, each with an alignment that has 17 congruent regions – 5 above the species level – and only 1 overlapping articulation. Two of the 12 scenarios (scenarios 1 and 21) that constitute pattern (3) are inverse move scenarios; however, this has no effect on the alignment pattern (compare Figs. 8A and 8B).

**Fig. 7.**
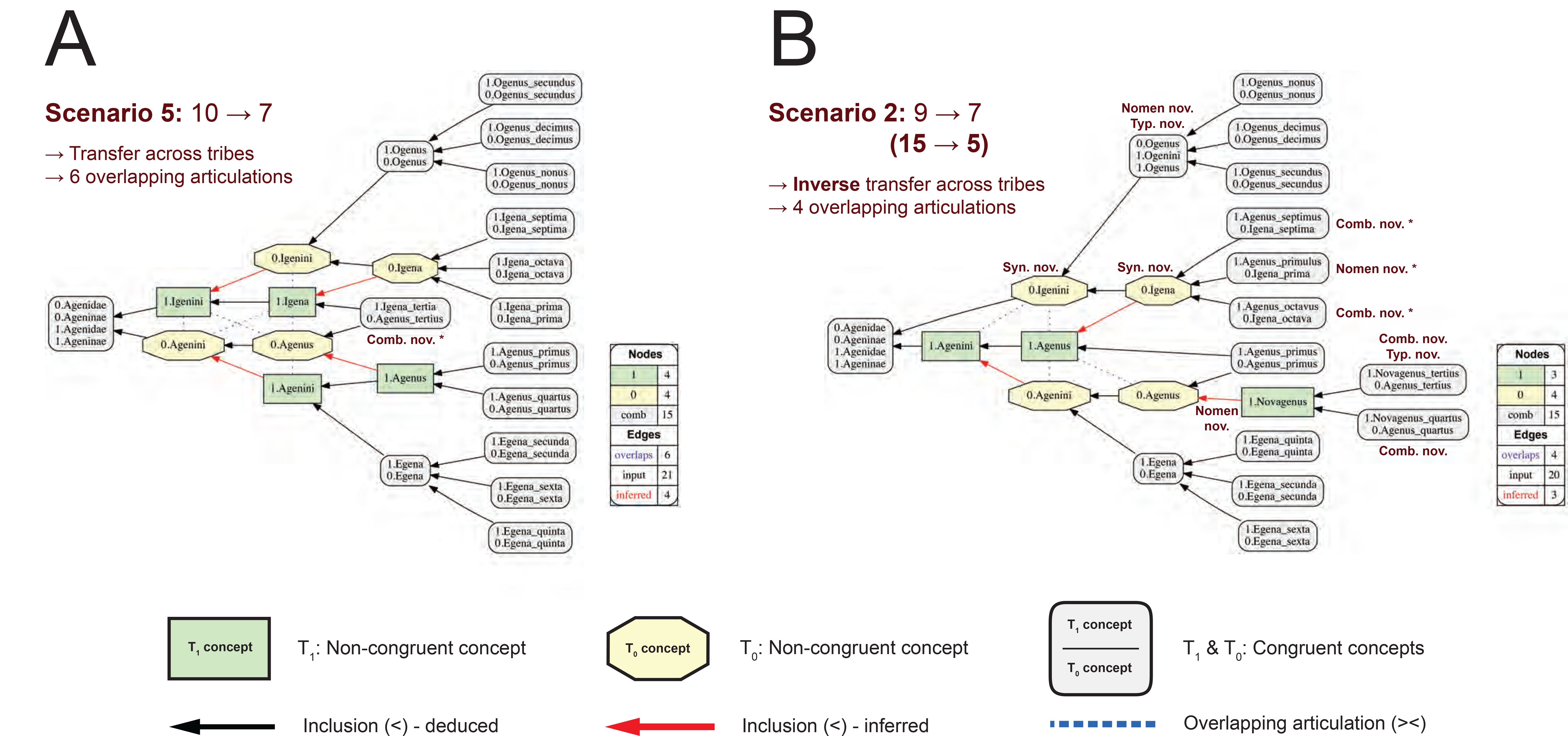

**Fig. 8.**
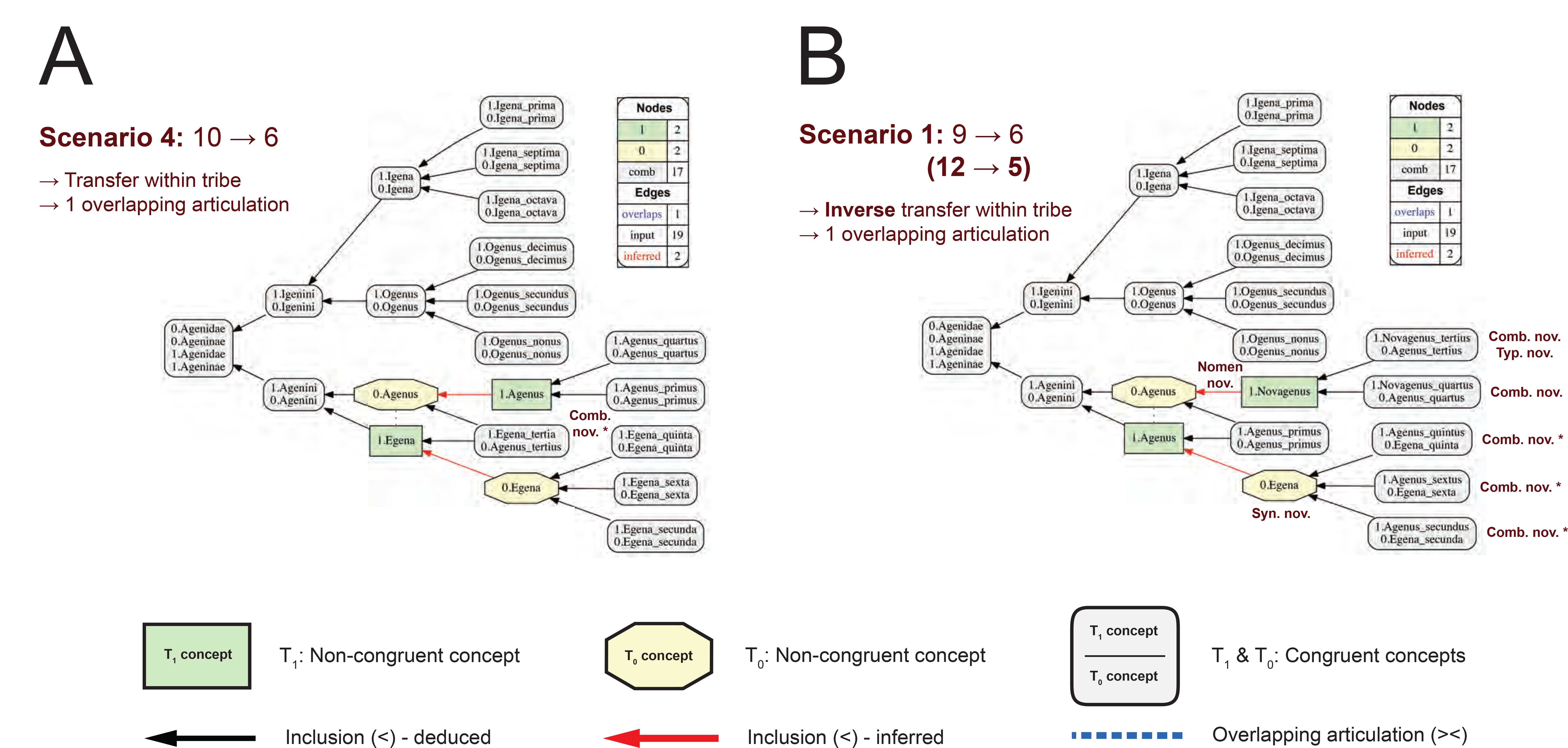

Comparison of the RCC-5 alignment representations and the nomenclatural change scenarios shows two rather different approaches to modeling identity and change relationships for this use case (Table 3, last column). In particular, patterns (1) and (2) are by some measure the most taxonomically impacting: the reinterpretation of a type specimen triggers that a species-level entity is transferred from one tribe-level entity to another (as conceived at t = 0). This move consistently translates into 6 or 4 overlapping articulations among the affected genus- and tribe-level concepts, depending on whether it is carried out in the intended or inverse direction. Nomenclaturally, this pattern does not hold well (compare Figs. 7A and 7B). Pattern (1) is variously associated with 1-11 nomenclatural changes (average: 2.9 ± 3.0), whereas pattern (2) involves 8-14 nomenclatural changes (average: 11.0 ± 2.4). The nomenclatural change trend between patterns (1) and (2) (less → more) is somewhat opposed to that of the RCC-5 alignments (more → less), in addition to being less homogenous. The greater average number of changes under pattern (2) is caused by global Priority constraints requiring inverse moves. Pattern (3) is taxonomically less impacting in comparison, because the reinterpretation of the type and associated species-level node transfer are confined to one tribe-level entity (as recognized at t = 0). Nevertheless, this relatively minor reclassification translates into a range of 2-11 nomenclatural changes (average: 4.3 ± 3.7), with either 2 or 7 changes in transfer scenarios in the intended direction, and 11 changes in transfer scenarios in the inverse direction (compare Figs. 8A and 8B).

## Discussion

### Compatibility of Nomenclature and Computational Logic

We have shown, likely to an unprecedented degree of sophistication, that many overarching principles of zoological nomenclature can be modeled with computational logic. Such logic can then be applied to specific taxonomic change scenarios to yield diverse but nomenclaturally accurate reasoning outcomes. While our approach and encoded program are not immediately designed to resolve real-life reclassification challenges, they demonstrate that ASP representation and reasoning powers are fully adequate to do so. In short, the rules of zoological nomenclature are logically tractable.

We reach this conclusion not because we have covered every principle, application, and exception represented in the 90 articles of the ICZN (1999). For instance, our use case does not account for the principle of the First Reviser (Article 24) or for the use of the Commission’s Plenary Power (Article 81) to preserve or suppress names or other nomenclatural acts under certain conditions, and typically with the intent to conserve “prevailing usage” (Vences et al. 2013). We have also not addressed issues of uncertainty in taxonomic judgment (Confalonieri et al. 2012). However, in the language of ASP these are simply additional constraints, each with specific trigger conditions (some of which may require user interaction). As needed, each novel constraint can be added to the elaboration-tolerant code to model increasingly complex cases.The size of the input taxonomies can be augmented, and additional variables such as author names or page numbers can be included to resolve Homonymy or Priority disputes more finely. Constraints can be applied to specific taxonomic ranks (“groups”). More input classifications and more steps can be accommodated.

We see no elements in the ICZN (1999) that are principally incompatible with ASP representation and reasoning to produce valid stable models for complex nomenclatural/taxonomic transition scenarios. Once the appropriate rules are encoded, the burden of producing viable stable models will depend largely on the systematists’ ability to specify sufficiently precise starting conditions upon which these rules can act.

However, demonstrating the ability to model nomenclatural change in logic is not the same as showing that such an approach is worth the effort. Hence we should ask: under what circumstances might the benefits of applying ASP logic outweigh the costs represented by encoding the rules, starting conditions, and then selecting suitable answer sets? The best but nevertheless tentative answer we have points to global nomenclatural registry and vetting services. Virtual environments such as ZooBank (Pyle and Michel 2008; Penev et al. 2016; Pyle 2016) aspire to act as globally coordinated repositories for taxonomic and nomenclatural acts.For instance, since 2012 registration with ZooBank is mandatory for electronic publications to acquire availability under ICZN rules (Krell and Pape 2015).

In the case of publishing complex sets of nomenclatural acts – such as those under consideration for the fruit fly name *Drosophila* where approximately 1500 valid names are in play (van der Linde and Houle 2008; O’Grady and Markow 2009; ICZN 2010) – ASP-encoded logic could be used to explore alternative valid change scenarios. Given multiple nomenclaturally viable options, the application logic could be further optimized identity solutions that maximize stability under certain prioritized criteria. Such criteria might include (1) minimize the total number of nomenclatural changes, or (2) minimize the number of changes at the generic level, or (3) maximize stability for the most frequently used binomials, etc. A likely less strict set of rules could nevertheless be applied for higher-level names where the Codes have no jurisdiction (Dubois 2015), and strive to maximize naming stability in light of taxonomic change at these supra-familiar levels.

In addition, ASP logic could be used as a means of verifying compliance of proposed emendations with nomenclatural rules, by blocking new submissions to the registry that are not consistent with any stable model. Following the rules of the ICZN (1999) is not always trivial, even for systematist users, and hence an automated service that checks for logic errors in human-proposed nomenclatural emendations could increase user confidence in enacting necessary changes and reduce rule-violating errors.

We realize that the suggested applications of ASP logic to nomenclatural registries require much further development – conceptual, technical, and social – to possibly lead to real-life implementations. This study is limited to illustrating the general compatibility of zoological nomenclature and computational logic. Yet we may also remind ourselves that the enterprise of discovering and naming nature’s diversity is far from finished. While estimates of global species richness remain uncertain, most indicate that millions of such entities remain unnamed (Caley et al. 2014). Future additions, synonymizations, and rearrangements of entities at varying taxonomic levels may have minimal or rather dramatic nomenclatural consequences. We hope that the scale of this challenge will motivate further work aimed at developing logic-enabled solutions based on this foundation-laying study.

### Computational Logic as an Assessment Tool for Nomenclatural Constraints

Modeling nomenclatural change scenarios in ASP logic may also give us a new perspective on the nature of nomenclature itself. In particular, our contrived use case illustrates how Priority and Coordination constraints may create dissimilar nomenclatural outcomes for what could be regarded as taxonomically identical, or at least highly comparative, triggers of change. From a strictly taxonomic perspective, the reinterpretation of the type specimen of *A. primus* and *A. tertius* as having a hemi-elliptical shaped instead of a square shape (as originally assessed), may by viewed as ‘the same amount of taxonomic change’. In each case, it would appear in retrospect that a single, species-level entity had been misdiagnosed and hence taxonomically misplaced. However, Priority makes these two scenarios highly unequal, considering that *A. primus* was coined in 1775 and *A. tertius* was coined in 1950 (Fig. 1A). Consequently, the reinterpretation of the older type ‘costs’ 14 nomenclatural emendations (Figs. 3B and 7A), whereas that of the younger type ‘costs’ 2 such changes (Figs. 1C and 7A).

Generally speaking, it appears that zoological nomenclature is designed so as to make changes to the taxonomic identities of names with older ages more costly, whereas those of younger names are less costly. Clearly such a design achieves that there is increased stability in *naming* taxonomic entities. Older names tend to persist in part by virtue of their higher age. But this is not necessarily the same as optimizing change ratios for the *interaction* of nomenclature and taxonomy. For that to be the case, one might instead design a naming system whose behavior in light of taxonomic change is less driven by Priority – which along with Coordination can produce highly unequal cascading effects – and more receptive to the ‘relative amount’ of taxonomic change to be enacted.

The RCC-5 alignments shown in Figs. 7 and 8 (see also Table 3) are aiming in this direction, and mitigate the more dramatic nomenclatural disparities to up to point, while still also modeling valid nomenclatural conditions for the starting and ending conditions. In particular, one might argue that type specimen reinterpretations that motivate taxonomic rearrangements of species-level entities spanning across multiple tribes (Fig. 7) should be consistently more costly than within-tribe moves (Fig. 8). After all, alignments for the former scenarios only generate 15 congruent Euler regions, whereas the latter generate 17 congruent Euler regions. However, such considerations of the relative *taxonomic* quantity of change are not primary design features of the ICZN (1999) principles and rules.

The history of Codes is deep and multi-faceted (Winsor 2001; Minelli 2003; Schuh 2003; Dayrat 2010; Dubois 2011; David et al. 2012). For instance, Witteveen (2016) connects the origins or the modern type concept in nomenclature to the struggle between the “metropolitan establishment” and “provincial radicals” in the colonial 19^th^ century United Kingdom. This struggle to avert the “chaos of synonymy” is ultimately grounded in human cognitive constraints. Atran (1998) argues that the Linnaean naming tradition retains important features of folk biology, and in this sense is well aligned what he calls cognitive universals. These include evolutionarily constrained notions of taxonomic rank, generic species, and rank-associated essences. Both folk biology, which is functional in localized ecological contexts, and Linnaean classification, which is global in scope, are designed to satisfy our cognitive preferences to maximize our inductive learning and reasoning potential for the given context.

Our logic explorations show that the principles of nomenclature are not designed in the main to respond *proportionally* to complex nomenclatural and taxonomic change scenarios. The principles may perform best under the cognitive premise that old and long-established names experience relatively minimal taxonomic reassignments at late stages in the history of human taxonomic making. If on the other hand the taxonomic identities of these early-period names are frequently redefined ‘late in the game’, then this may result in disproportionately many nomenclatural emendations. Another way of saying this is that the Linnaean system may be biased to favor early success in achieving natural, inductively projectible classifications. Such early success is certainly possible if perceived taxonomic groups are not very diverse and are otherwise favorably accessible to human cognition. However, other complex groups may well experience abundant and significant late-stage reassessments – a circumstance that in some sense runs counter to both our cognitive biases and those (logically) embedded in the Codes of nomenclature. Many naming controversies in systematics reflect the tension of apparent early-versus late-stage success in recognizing complex natural entities and relationships.

Applications of computational logic to nomenclatural/taxonomic change scenarios can inform future designs of identifier systems for systematics that find the best balance between human cognitive preferences and logic-informed representation and reasoning maxims. We have shown that the ASP approach can provide both diagnostic and prescriptive input in this regard. It is not too late do pursue this path, and doing so may enable us to more fully bring nomenclature into the realm of computationally enabled big data science (Page 2016).

## Acknowledgments

The authors thank Stanley Blum, Neal Evenhuis, Thomas Pape, David Patterson, Richard Pyle, and Francisco Welter-Schultes for their advice in conceiving the use case and in the application of nomenclatural rules. Bertram Ludäscher, Parisa Kianmajd, and Shizuo Yu provide support for the Euler/X toolkit.

## Supplementary Materials

Supplementary Materials S1. – Answer Set Programming code (.txt), written in F2LP and with extensive comments, to perform the 20 nomenclatural taxon use case in conjunction with the Potassco solver *clingo*. See also Table 1.

Supplementary Materials S2. – Data file (.txt) with installation instructions and command line interface operations to run the ASP code and obtain the related reasoning and visualization outputs.

Supplementary Materials S3. – Data file (.txt) with the complete textual *clingo* solver output for the 20 nomenclatural taxon use case. This output is used to produce the GraphViz visualizations.

Supplementary Materials S4. – Java Archive file (.jar) needed to translate the *clingo* solver output into the corresponding GraphViz visualizations.

Supplementary Materials S5. – Collated set of GraphViz visualizations (transformed into.pdf) for the 20 nomenclatural taxon use case.

Supplementary Materials S6. – Set of 36 Euler input data files for the respective alignments (summarized in Table 3). Each file is saved is saved in .txt format and contains annotations and instructions for run commands to yield the alignments and input/output visualizations.

Supplementary Materials S7. – Set of 36 Euler/X toolkit output *Maximally Informative Relations* (MIR) for the input data files provided in the Supplementary Materials S6. Each output file is saved in .csv format.

Supplementary Materials S8. – Set of 36 Euler/X output alignment visualizations (.pdf) for the input data files provided in the Supplementary Materials S6.

## References

Atran S. 1998. Folk biology and the anthropology of science:cognitive universals and cultural particulars. Behav. Brain Sci. 21:547–569.

Baader F., Horrocks I., Sattler U. 2008. Description logics. In: van Harmelen F., Lifschitz V., Porter B., editors. Handbook of Knowledge Representation. Amsterdam: Elsevier Academic Press. p. 135–179.

Brachman R.J., Levesque, H.J. 2004. Knowledge representation and reasoning. San Francisco, CA: Morgan Kaufmann Publishers.

Brewka G., Eiter T., Truszczyński M. 2011. Answer set programming at a glance. Commun. ACM 54:92–103.

Brinckmann-Voss A., Calder D.R. 2013. Zyzzyzus rubusidaeus (Cnidaria, Hydrozoa, Tubulariidae), a new species of anthoathecate hydroid from the coast of British Columbia, Canada. Zootaxa 3666:389–397.

Brooks D.R., Erdem E., Erdogan S.T., Minett J.W., Ringe D. 2007. Inferring phylogenetic trees using Answer Set Programming. J. Autom. Reasoning 39:471–511.

Bryant H.N., Cantino P.D. 2002. A review of criticisms of phylogenetic nomenclature:is taxonomic freedom the fundamental issue? Biol. Rev. 77:39–55.

Caley M.J., Fisher R., Mengersen K. 2014. Global species richness estimates have not converged. Trends Ecol. Evol. 29:187–188.

Cappellini E., Gentry A., Palkopoulou E., Ishida Y., Cram D., Roos A.-M., Watson M., Johansson U.S., Fernholm B., Agnelli P., Barbagli F., Littlewood D.T.J., Kelstrup C.D., Olsen J.V., Lister A.M., Roca A.L., Dalén L., Gilbert M.T.P. 2013. Resolution of the type material of the Asian elephant, Elephas maximus Linnaeus, 1758 (Proboscidea, Elephantidae). Zool. J. Linn. Soc. 170:222–232.

Casadio C., Lambek J. 2005. A computational algebraic approach to Latin grammar. Res. Lang. Comput. 3:45–60.

Chawuthai R., Takeda H., Wuwongse V., Jinbo U. 2013. A logical model for taxonomic concepts for expanding knowledge using Linked Open Data. In: Larmande P., Arnaud E., Mougenot I., Jonquet C., Libourel T., Ruiz M., editors. S4BioDiv 2013, Semantics for Biodiversity – Proceedings of the First International Workshop on Semantics for Biodiversity, Montpellier, France, May 27, 2013. CEUR Workshop Proceedings, Vol-797. p. 1–8.

Chen M., Yu S., Franz N., Bowers S., Ludäscher B. 2014. Euler/X:a toolkit for logic-based taxonomy integration. arXiv:1402.1992 [cs.LO] Available at Accessed June 15, 2016.

Confalonieri R., Nieves J.C., Osorio M., Vázquez-Salceda J. 2012. Dealing with explicit preferences and uncertainty in answer set programming. Ann. Math. Artif. Intell. 65:159–198.

David J., Garrity G.M., Greuter W., Hawksworth D.L., Jahn R., Kirk P.M., McNeill J., Michel E., Knapp S., Patterson D.J., Tindall B.J., Todd J.A., Tol J., Turland N.J. 2012. Biological nomenclature terms for facilitating communication in the naming of organisms. ZooKeys 192:67–72.

Dayrat B. 2010. Celebrating 250 dynamic years of nomenclatural debates. In: Polaszek A., editor. Systema Naturae 250 – The Linnaean Ark. Boca Raton, FL: CRC Press p. 186–239.

Deans A.R., Lewis S.E., Huala E., Anzaldo S.S., Ashburner M., Balhoff J.P., Blackburn D.C., Blake J.A., Burleigh J.G., Chanet B., Cooper L.D., Courtot M., Csösz S., Cui H., Dahdul W., Das S., Dececchi T.A., Dettai A., Diogo R., Druzinsky R.E., Dumontier M., Franz N.M., Friedrich F., Gkoutos G.V., Haendel M., Harmon L.J., Hayamizu T.F., He Y., Hines H.M., Ibrahim N., Jackson L.M., Jaiswal P., James-Zorn C., Köhler S., Lecointre G., Lapp H., Lawrence C.J., Le Novère N., Lundberg J.G., Macklin J., Mast A.R., Midford P.E., Mikó I., Mungall C.J., Oellrich A., Osumi-Sutherland D., Parkinson H., Ramfrez M.J., Richter S., Robinson R.N., Ruttenberg A., Schulz K.S., Segerdell E., Seltmann K.C., Sharkey M.J., Smith A.D., Smith B., Specht C.D., Squires R.B., Thacker R.W., Thessen A.E., Fernandez-Triana J., Vihinen M., Vize P.D., Vogt L., Wall C.E., Walls R.L., Westerfeld M., Wharton R.A., Wirkner C.S., Woolley J.B., Yoder M.J., Zorn A.M., Mabee P. 2015. Finding our way through phenotypes. PLoS Biol. 13:e1002033.

Dmitriev D.A., and Yoder M. 2016. NOMEN – a nomenclatural ontology for biological names (not concepts). Available at Accessed June 15, 2016.

Donnellan KS. 1966. Reference and definite descriptions. Phil. Review 77:281–304.

Dubois A. 2011. The International Code of Zoological Nomenclature must be drastically improved before it is too late. Bionomina 2:1–104.

Dubois A. 2015. The Duplostensional Nomenclatural System for higher zoological nomenclature. Dumerilia 5:1–108.

Eiter T. 2008. Answer Set Programming in a nutshell. Available at gradlog.informatik.uni-freiburg.de/gradlog/slides_ak/eiter_asp.pdf Accessed June 15,2016.

Eiter T., Ianni G., Krennwallner T. 2009. Answer Set Programming:a primer. In: Tessaris S., Franconi E., Eiter T., Gutierrez S., Handschuh S., Rouset M.-C., Schmidt R.A., editors. Reasoning Web. Semantic Technologies for Information Systems: 5th International Summer School 2009, Brixen-Bressanone, Italy, August 30 – September 4, 2009, Tutorial Lectures.Lect. Notes Comput. Sc. 5689:40–110.

Eiter T., Ianni G., Lukasiewicz T., Schindlauer R., Tompits H. 2008. Combining answer set programming with description logics for the Semantic Web. Artif. Intell. 172:1495–1539.

Ereshefsky M. 2007. Foundational issues concerning taxa and taxon names. Syst. Biol. 56:295–301.

Franz N.M. 2005. On the lack of good scientific reasons for the growing phylogeny/classification gap. Cladistics 21:495–500.

Franz N.M., Chen M., Kianmajd P., Yu S., Bowers S., Weakley A.S., Ludäscher B. 2016a. Names are not good enough:reasoning over taxonomic change in the Andropogon complex. Semantic Web 7:1–23. (Early Access) DOI:10.3233/SW-160220

Franz N.M., Chen M., Yu S., Kianmajd P., Bowers S., Ludäscher B. 2015. Reasoning over taxonomic change:exploring alignments for the Perelleschus use case. PLoS ONE 10(2):e0118247.

Franz N.M., Peet R.K., Weakley A.S. 2008. On the use of taxonomic concepts in support of biodiversity research and taxonomy. In: Wheeler Q.D., editor. The New Taxonomy,Systematics Association Special Volume Series 74. Boca Raton, FL: Taylor & Francis. p. 63–86.

Franz N.M, Pier N.M., Reeder D.M., Chen M., Yu S., Kianmajd P., Bowers S., Ludäscher B. 2016b. Two influential primate classifications logically aligned. Syst. Biol. 65:1–22. (Early Access) DOI:10.1093/sysbio/syw023.

Franz N.M., Thau D. 2010. Biological taxonomy and ontology development:scope and limitations. Biodiv. Informatics 7:45–66.

Gansner ER, North SC. 2000. An open graph visualization system and its applications to software engineering. Softw. Pract. Exp. 30:1203–1233.

Gebser M., Kaminski R., Kaufmann B., Lindauer M., Ostrowski M., Romero J., Schaub T., Thiele S. 2015. Potassco user guide. Available at sourceforge.net/projects/potassco/files/potassco_guide/ Accessed June 15, 2016.

Gebser M., Kaufmann B., Kaminski R., Ostrowski M., Schaub T., Schneider M.T. 2011. Potassco:the Potsdam Answer Set Solving Collection. AI Commun. 24:107–124.

Gebser M., Schaub T., Thiele S., Usadel B., Veber P. 2008. Detecting inconsistencies in large biological networks with Answer Set Programming. In: de la Banda M.G., Pontelli E., editors. Logic Programming. Proceedings of the 24^th^ International Conference, ICLP 2008, Udine, Italy, December 9-13. Lect. Notes Comput. Sc. 5366:130–144.

Gelfond M. 2008. Answer sets. In: van Harmelen F., Lifschitz V., Porter B., editors. Handbook of Knowledge Representation. Amsterdam: Elsevier Academic Press. p. 285–316.

Gelfond M., Lifschitz V. 1988. The Stable Model Semantics For Logic Programming. In: Kowalski R.A., Bowen K.A., editors. Logic Programming, Proceedings of the Fifth International Conference and Symposium, Seattle, Washington, August 15-19, 1988 (2 Vols). Cambridge: MIT Press. p. 1070–1080.

Grenon P., Smith B., Goldberg L. 2004. Biodynamic ontology:applying BFO in the biomedical domain. Stud. Health. Technol. Inform. 102:20–38.

Hawksworth D.L. 2001. Nomenclature, systems.In: Levin S.A., editor. Encyclopedia of Biodiversity, Vol. 4. Amsterdam: Elsevier Academic Press. p. 389–402.

ICZN – International Commission on Zoological Nomenclature. 1999. International Code of Zoological Nomenclature. Fourth Edition. London, UK:International Trust for Zoological Nomenclature.

ICZN – International Commission on Zoological Nomenclature. 2010. Opinion 2245 (Case 3407) Drosophila Fallén, 1823 (Insecta, Diptera): Drosophila funebris Fabricius, 1787 is maintained as the type species. Bull. Zool. Nom. 76:106–115.

Krell F.-T., Pape T. 2015.Electronic publications need registration in ZooBank to be available. Bull. Zool. Nomencl. 72:245–251.

Kutz O., Mossakowski T., Lücke D. 2010. Carnap, Goguen, and the hyperontologies: logical pluralism and heterogeneous structuring in ontology design. Log. Univers. 4:255–333.

Laloy F., Rage J.-C., Evans S.E., Boistel R., Lenoir N., Laurin M. 2013. A re-interpretation of the Eocene anuran Thaumastosaurus based on MicroCT examination of a ‘mummified’ specimen. PLoS ONE 8(9):e74874.

Lee J., Palla R. 2009. System F2LP – computing Answer Sets of First-Order formulas. In: Erdem E., Lin F., Schaub T., editors. Proceedings Of The 10Th International Conference On Logic Programming And Nonmonotoning Reasoning (Lpnmr 2009), Lpnmr 2009, Potsdam, Germany. Lect. Notes Comp. Sci. 5753:515–521.

Lifschitz H. 2008. What is answer set programming? In: Proceedings of the 23rd AAAI Press Conference on Artificial Intelligence, AAAI 2008, Menlo Park, CA, USA. p. 1594–1597.

Lifschitz V., Turner H. 1999. Representing transition systems by logic programs. In: Gelfond M., Leone N., Pfeifer G., editors. Proceedings of the Fifth International Conference on Logic Programming and Nonmonotonic Reasoning (LPNMR-99), El Paso, Texas. Lect. Notes Artif. Int. 1730:92–106.

Linnaeus C. 1753. Species plantarum: exhibentes plantas rite cognitas, ad genera relatas, cum differentiis specificis, nominibus trivialibus, synonymis selectis, locis natalibus, secundum systema sexuale digestas; 2 vols. Holmiae: Impensis Laurentii Salvii.

Linnaeus C. 1758. Systema nature per regna tria nature, secundum classes, ordines, genera, species, cum characteribus, differentiis, synonymis, locis. Tomus I & II. Editio decima, reformata. Holmis: Salvius.

McCarthy J. 1998. Elaboration tolerance. In: Symposium on Logical Formalizations of Commonsense Reasoning (Common Sense 98), January 07-09, 1998, University of London, London, UK. p. 198–216.

McNeill J., Turland N.J., Barrie F.R., Buck W.R., Greuter W., Wiersema J.H. 2012. International Code of Nomenclature for Algae, Fungi, and Plants (Melbourne Code). Königstein: Koeltz Scientific Books.

Midford P.E., Dececchi T.A., Balhoff J.P., Dahdul W.M., Ibrahim N., Lapp H., Lundberg J.G., Mabee P.M., Sereno P.C., Westerfield M., Vision T.J., Blackburn D.C. 2013. The vertebrate taxonomy ontology: a framework for reasoning across model organism and species phenotypes. J. Biomed. Semant. 4:34.

Minelli A. 2003. Historical Review Of Systematic Biology And Nomenclature. In: Contrafatto G., Minelli A., editors. Biological Science Fundamentals (Systematics), in Encyclopedia of Life Support Systems (EOLSS). Oxford, UK: Eolss Publishers. p. 1–13.

Mungall C.J., Gkoutos G.V., Smith C.L., Haendel M.A., Lewis S.E., Ashburner M. 2010. Integrating phenotype ontologies across multiple species Genome Biol. 2010, 11:R2.

Nicholson K.E., Crothier B.I., Guyer C., Savage J.M. 2012. It is time for a new classification of anoles (Squamata: Dactyloidae). Zootaxa 3477:1–108.

O’Grady P.M., Markow T.A. 2009. Phylogenetic taxonomy in Drosophila. Fly (Austin)3:10–14.Page R.D.M. 2016. Surfacing the deep data of taxonomy. ZooKeys 550:247–260.

Panahiazar M., Sheth A.P., Ranabahu A., Vos R.A., Leebens-Mack, J. 2013. Advancing data reuse in phyloinformatics using an ontology-driven Semantic Web approach. BMC Med. Genomics 2013, 6(Suppl. 3):S5.

Patterson D.J., Cooper J., Kirk P.M., Pyle R.L., Remsen D.P. 2010. Names are key to the big new biology. TREE 25:686–691.

Patterson D., Mozzherin D., Shorthouse D., Thessen A. 2016 Challenges with using names to link digital biodiversity information. Biodivers. Data J. 4:e8080.

Penev L., Paton A., Nicolson N., Kirk P., Pyle R.L., Whitton R., Georgiev T., Barker C., Hopkins C., Robert V., Biserkov J., Stoev P. 2016. A common registration-to-publication automated pipeline for nomenclatural acts for higher plants (International Plant Names Index, IPNI), fungi (Index Fungorum, MycoBank) and animals (ZooBank). In: Michel E., editor. Anchoring Biodiversity Information: from Sherborn to the 21^st^ Century and Beyond. ZooKeys 550:233–246.

Pyle R.L. 2016. Towards a Global Names Architecture:the future of indexing scientific names. In: Michel E., editor.Anchoring Biodiversity Information: from Sherborn to the 21^st^ Century and Beyond. ZooKeys 550:261–281.

Pyle R.L., Michel E. 2008. ZooBank:developing a nomenclatural tool for unifying 250 years of biological information. Zootaxa 1950:39–50.

Reiter R. 1980. A logic for default reasoning. Artif. Intell. 13:81–132.

Remsen D. 2016. The use and limits of scientific names in biological informatics. ZooKeys 550:207–223.

Rhodin A.G.J., Carr J.L. 2009. A quarter millenium of uses and misuses of the turtle name Testudo scabra: identification of the type specimens of T. scabra Linnaeus 1758 (= Rhinoclemmys punctularia) and T. scripta Thunberg in Schoepff 1792 (= Trachemys scripta scripta). Zootaxa 2226:1–18.

Ribot F., Gibert J., Harrison T. 1996. A reinterpretation of the taxonomy of Dryopithecus from Valles-Penedes, Catalonia (Spain). J. Hum. Evol. 31:129–141.

Schuh R.T. 2003. The Linnaean system and its 250-year persistence. Bot. Rev. 69:59–78.

Sereno P.C. 2005. The logical basis of phylogenetic taxonomy. Syst. Biol. 54:595–619.

Smith B., Ashburner M., Rosse C., Bard J., Bug W., Ceusters W., Goldberg L.J., Eilbeck K., Ireland A., Mungall C.J., The Obi Consortium, Leontis N., Rocca-Serra P., Ruttenberg A., Sansone S.-A., Scheuermann R.H., Shah N., Whetzel P.L., Lewis S. 2007. The OBO Foundry:coordinated evolution of ontologies to support biomedical data integration. Nature Biotech. 25:1251–1255.

Sterner B.W., Franz N.M. 2016. Cognitive pragmatics for big biodiversity data: taxonomy for humans or omputers? Biol. Theory. (In Review)

Thessen A.E., Bunker D.E., Buttigieg P.L., Cooper L.D., Dahdul W.M., Domisch S., Franz N.M., Jaiswal P., Lawrence-Dill C.J., Midford P.E., Mungall C.J., Ramírez M.J., Specht C.D., Vogt L., Vos R.A., Walls R.L., White J.W., Zhang G., Deans A.R., Huala E., Lewis S.E., Mabee P.M. 2015. Emerging semantics to link phenotype and environment. PeerJ 3:e1470.

Tuominen J., Laurenne N., Hyvönen E. 2011. Biological Names And Taxonomies On The Semantic Web – Managing The Change In Scientific Conception. In: Antoniou G., Grobelnik M., Simperl E., Parsia B., Plexousakis D., Leenheer P., Pan J., editors. The Semantic Web:Research and Applications. Lect. Notes Comput. Sc. 6644:255–269.

van der Linde K., Houle D. 2008. A supertree analysis and literature review of the genus Drosophila and closely related genera (Diptera, Drosophilidae). Insect Syst. Evol. 39:241–267.

Vane-Wright R.I. 2003. Indifferent philosophy versus almighty authority: on consistency, consensus and unitary taxonomy. Syst. Biodiv. 1:3–11.

Vences M., Guayasamin J.M., Miralles A., de la Riva, I. 2013. To name or not to name:criteria to promote economy of change in Linnaean classification schemes. Zootaxa 3636:201–244.

Walls R.L., Deck J., Guralnick R., Baskauf S., Beaman R., Blum S., Bowers S., Buttigieg P.L., Davies N., Endresen D., Gandolfo M.A., Hanner R., Janning A., Krishtalka L., Matsunaga, Midford P., Morrison N., Tuama É.Ó., Schildhauer M., Smith B., Stucky, B., Thomer A., Wieczorek J., Whitacre J., Wooley J. 2014. Semantics in support of biodiversity knowledge discovery: an introduction to the Biological Collections Ontology and related ontologies. PLoS ONE 9(3):e89606.

Winsor M.P. 2001. Cain on Linnaeus: the scientist-historian as unanalysed entity. Stud. Hist Phil. Biol. Biomed. Sci. 32:239–254.

Witteveen J. 2015. Naming and contingency:the type method of biological taxonomy. Biol.Phil. 30:569–586.

Witteveen J. 2016. Suppressing synonymy with a homonym:the emergence of the nomenclatural type concept in nineteenth century natural history. J. Hist. Biol. 49:135–189.

Woodley M.A., Naish D., McCormick C.A. 2011. A baby sea-serpent no more: reinterpreting Hagelund’s juvenile “cadborosaur” report. J. Sci. Explor. 25:495–512.

